# Cell proliferation and migration explain pore bridging dynamics in 3D printed scaffolds of different pore size

**DOI:** 10.1101/2020.03.12.989053

**Authors:** Pascal R. Buenzli, Matthew Lanaro, Cynthia S. Wong, Maximilian P. McLaughlin, Mark C. Allenby, Maria A. Woodruff, Matthew J. Simpson

**Affiliations:** School of Mathematical Sciences, Queensland University of Technology (QUT), Brisbane, Australia; School of Mechanical Medical and Process Engineering, Queensland University of Technology (QUT), Brisbane, Australia

**Keywords:** Cell migration, cell proliferation, 3D printing, biofabrication, parameter estimation, mathematical modelling

## Abstract

Tissue growth in bioscaffolds is influenced significantly by pore geometry, but how this geometric dependence emerges from dynamic cellular processes such as cell proliferation and cell migration remains poorly understood. Here we investigate the influence of pore size on the time required to bridge pores in thin 3D-printed scaffolds. Experimentally, new tissue infills the pores continually from their perimeter under strong curvature control, which leads the tissue front to round off with time. Despite the varied shapes assumed by the tissue during this evolution, we find that time to bridge a pore simply increases linearly with the overall pore size. To disentangle the biological influence of cell behaviour and the mechanistic influence of geometry in this experimental observation, we propose a simple reaction–diffusion model of tissue growth based on Porous-Fisher invasion of cells into the pores. First, this model provides a good qualitative representation of the evolution of the tissue; new tissue in the model grows at an effective rate that depends on the local curvature of the tissue substrate. Second, the model suggests that a linear dependence of bridging time with pore size arises due to geometric reasons alone, not to differences in cell behaviours across pores of different sizes. Our analysis suggests that tissue growth dynamics in these experimental constructs is dominated by mechanistic crowding effects that influence collective cell proliferation and migration processes, and that can be predicted by simple reaction–diffusion models of cells that have robust, consistent behaviours.

## 1 Introduction

The traditional cell culture on flat surfaces or in liquids is transitioning in favour of bio-mimicking porous 3D scaffolds to provide realistic substrates for cell therapies and disease models [1–5] Although 3D scaffolds can approximate the form and function of natural extracellular matrices, many remain limited due to inconsistent scaffold architecture. Recent 3D printing (3DP) technologies include melt electrowriting that enable precise microscale manufacture of cell culture scaffolds [6–9]. This consistency could allow a more complete control over cell behaviours, such as controlling cell proliferation and cell migration. However, explicit quantitative correlations between cell behaviour and scaffold architecture have yet to be identified. Mathematical relationships could define robust design protocols for the emerging tissue biomanufacturing industry [10, 11].

There are many design questions that need to be addressed in the production of these scaffolds, such as determining their optimal size, shape and material properties [2–4]. These properties have been shown to impact cell attachment, viability, proliferation, migration, and differentiation, among other functions [12–18], and they could be tuned to control the manufacture of complex multicellular tissues or organs. Recent additive manufacturing techniques have leveraged these biophysical relationships in an ad hoc manner: by 3D printing bilayer cylindrical scaffolds as vascular grafts with endothelial and muscle cells [19] or by patterning scaffold pores or fibres to spatially control cell morphology and differentiation [20]. Therefore, a predictive mathematical model of cell behaviour across 3D printing parameters could provide control or optimisation strategies to develop complex yet robust tissue biomanufacturing [11].

In this work, we focus on pore bridging experiments where quasi-two-dimensional tissue sheets are grow are grown in 3D-printed scaffolds by osteoblastic cells seeded onto the perimeter of square-shaped pores of different sizes. As the experiment proceeds, cells migrate and proliferate to form new tissue that extends inwards to eventually close or *bridge* the pores to form a sheet-like tissue structure. We use this experimental design to investigate two questions of central interest: (i) whether there is an optimal scaffold pore size *L* for the production of such tissues; and (ii) whether different choices of *L* lead to different cell-level mechanisms, e.g. different rates of cell migration and different rates cell proliferation. Our experimental design involves working with 3D pores with a simplified geometry. The vertical length scale of the pores (100 μm) is small compared to the horizontal length scale (200–600 μm). This simplified geometry allows us to describe the experimental data using appropriate 2D quantities, such as the pore area as a function of time. This design also alleviates the influence of 3D effects of tissue growth on the apparent, within-plane rate of growth of the tissue sheets, an influence that has been observed in scaffolds of large thickness (2000–6000 μm) [21–23]. Further, this simplified geometry allows us to work with a 2D mathematical modelling framework that describes cell density as a function of 2D position, *(x, y*), rather than a full 3D coordinate system, (x, *y, z*).

Previous work suggests that pore size could lead to differences in cellular behaviour, such as differences in the rate of cell migration and the rate cell proliferation. Cellular tensile stress correlates with cell proliferation in a number of experimental situations, including in tissue engineering and wound healing [12, 24, 25]. At the tissue interface, contractile actomyosin rings induce a tissue surface tension [26] subject to a curvature dependence similar to that of surface tension in fluids [13, 14, 17]. It is therefore reasonable to expect that tissue curvature influences cell proliferation and thus tissue growth. The crowding and spreading of tissue material near concavities and convexities of bone tissue was also found to involve curvature-dependent cell behaviours [27]. Curvature, in turn, is affected by the overall size of a pore: the larger a pore is, the smaller the average curvature. Mathematically, the average curvature of a two-dimensional pore cross section is always *2π/P*, where *P* is the perimeter of the pore.^1^ For square pores of side length *L*, we have *P* = 4L, so we may expect the rate of cell proliferation could depend on pore size *L* due to curvature. While previous studies have considered tissue growth in pores of different shapes [13, 21, 23], they did not systematically examine a broad range of pore sizes.

Experimental images of the cell bridging experiments indicate that varying the initial pore size *L* leads to visually distinct curvatures along the leading edge of the population of cells. Small pores lead to more rapid bridging than large pores, and those pores with smaller size tend to involve more rapid development of high curvature. However, it is unclear from the experimental data how much of the difference in bridging time is due to geometric reasons, and how much is due to differences in the proliferative and migratory behaviours of the cells. To gain insights into these questions, we propose a mathematical model that captures the mechanistic, geometric processes of tissue growth, and that leaves open for determination cell behavioural aspects. Comparing the experimental data and the model then allows us to determine important properties of these cell behavioural aspects.

Several mathematical modelling strategies have been developed to explore the interplay between tissue growth and curvature effects, which is an important area of research toward the industrial implementation of scaffold design optimisation [11], see Callens et al. [18] for a recent review. Mathematical models range from continuum mechanics growth laws [29–31, 5], computational models based on the idea of mean curvature flow, that borrow ideas from fluid mechanics and the role of surface tension but that do not consider cells explicitly [13, 14, 21–23, 15, 16, 32, 17, 33, 34], to other modelling approaches that consider the effects of cell-level behaviour, such as particle-based models with mechanical interaction and contractile forces at tissue interfaces [35], and models including tissue crowding and spreading during surface growth, which lead to hyperbolic curvature flow models [36, 27, 37].

In this work we propose a simple reaction–diffusion model, called the *Porous-Fisher* model [38], to describe the combined cell proliferation and cell migration that leads to new tissue formation in the scaffold pores. The Porous-Fisher model is an extension of the classical Fisher–Kolmogorov model which describes how a population of cells spreads spatially through the combined effects of cell migration and cell proliferation [39–41]. In the Fisher–Kolmogorov model, cell proliferation is assumed to follow a classical logistic growth model where the per capita growth rate of cells is a linearly decreasing function of density to simulate contact inhibition of proliferation [38, 42, 43]. Cell migration in the Fisher–Kolmogorov model is modelled by making the simplest assumption that cells migrate randomly. Therefore, the migration of cells is represented by a linear diffusion process where cell-to-cell interactions have no impact upon cell migration [38, 42, 43]. The Porous-Fisher model generalises linear diffusion to density-dependent diffusion, which accounts for cell-to-cell interactions. The density dependence of proliferation and migration in the Porous-Fisher model we consider represents a mechanistic influence of space constraints, i.e., the availability of space for cell motion and cell division, while per-capita parameters associated with these processes represent the cell behavioural aspects. This model has been widely used to model wound healing processes in two-dimensional scratch assays [42–48], as well as the outward growth of initially-confined populations of cells [49–51].

Unlike other studies that connect experimental observations with the Porous-Fisher model through counting cells and constructing cell density profiles [52], here we aim to use the model in a more practical way by connecting its outputs with very simple experimental observations, such as the time to bridge. We find that even this simple measurement provides very insightful mechanistic insight as to the proliferative and diffusive behaviour of the cells in pores of different sizes.

## 2 Materials and Methods

### 2.1 Tissue growth experiments

The full experimental protocol is described in [53]. In brief, polycaprolactone scaffolds were fabricated by melt electrospinning depositing fibres 50 μm in diameter [54], to manufacture scaffolds of size 7 mm×7 mm× 100 μm (2 fibre layers thick) with square-shaped pores of lengths ranging from *L* = 200 μm to *L* = 600 μm (Figure 1). A minimum of 5 × 5 square shaped pores were produced to omit any culture handling effects on the scaffold edges. The scaffolds were sterilised and incubated at 37°C in 5% CO_2_ and 95% air, in media overnight, prior to seeding.

**Figure 1.**
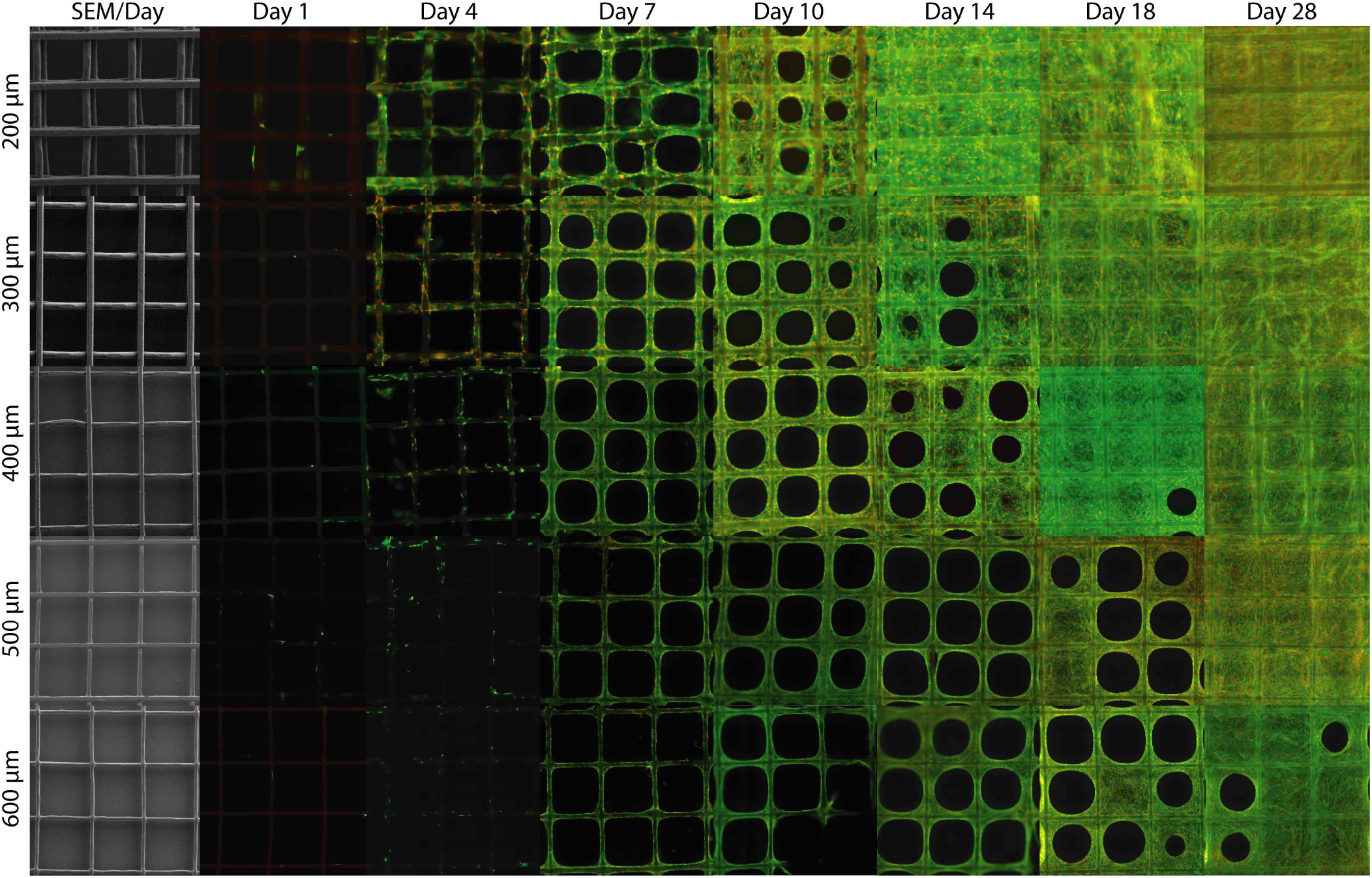
(Color online.) Experimental pore bridging in 3D printed scaffolds of different initial sizes (rows). Each time snapshot comes from a different experiment. The snapshots are rescaled to fit the central 3 × 3 pores of a larger scaffold into a fixed size. The first column shows initial scaffolds imaged by Scanning Electron Microscopy. The other images depict cell nuclei (red), cytoskeleton (green), and the remaining pore to bridge (black).

Prior to seeding, murine calvarial osteoblastic cells (MC3T3-E1) cells [55] were cultured in α-MEM, 10% fetal bovine serum, and 1% penicillin-streptomycin (Thermo Fisher). Cells were detached using 0.05% trypsin, manually counted and seeded at 7500 cells in 250 μL media onto each scaffold within a 48-well plate (Nunc, Thermo Fisher) on top of nonadherent 2% agarose to prevent cell-to-plate attachment. Cells were allowed to attach to the 3D printed scaffold for 4 h before adding the remaining 250 μL of media. Cell-seeded scaffolds were cultured in a humidified environment at 37 °C in 5% CO2 for 28 days. Media was changed every 2–3 days from day 1 to 14, every 1-2 days from day 15 to 21, then every 24 h from day 22 to 28. Cell viability was assessed at day 10, 14 and 28 using calcein AM and ethidium homodimer which stained live and dead cells, respectively [53]. Cell-seeded scaffolds were fixed in 4% paraformaldehyde at Day 1, 4, 7, 10, 14, 18, and 28 and tissue growth was assessed by staining cell nuclei with DAPI and actin filament with Alexa Fluor 488 phalloidin (Thermo Fisher). Cell and tissue morphology was visualised using fluorescent microscopy (Leica AF6000 LX) [53]. Each imaging measurement was repeated across *N* = 3 replicate scaffolds for each value of *L* = 200; 300; 400; 500 and 600μm, except Day 18 for *L* = 200; 400; and 500μm (*N* = 2), and measured on the central *n* = 9 pores per replicate scaffold.

From the microscope images (Figure 1) we estimated the time to bridge *T*_b_ for a given mesh size *L* by fitting the experimental time course of pore area normalised by initial pore area [53], denoted by *A(t)/A*(0), with a curve

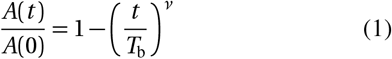

(see Figure 2 and supplementary Figure S1). Only data points before bridging occurs were considered for the fit, since *A*(*t*) ≡ 0 after bridging. Parameters of best fit *T*_b_ and *v* were found by minimising reduced *χ*^2^ (chi-squared) statistics using the nonlinear least-square Marquardt–Levenberg algorithm [56]. An estimate of the standard error on these fitted parameters is provided by the so-called asymptotic standard error during the fitting procedure [57, 56].

**Figure 2.**
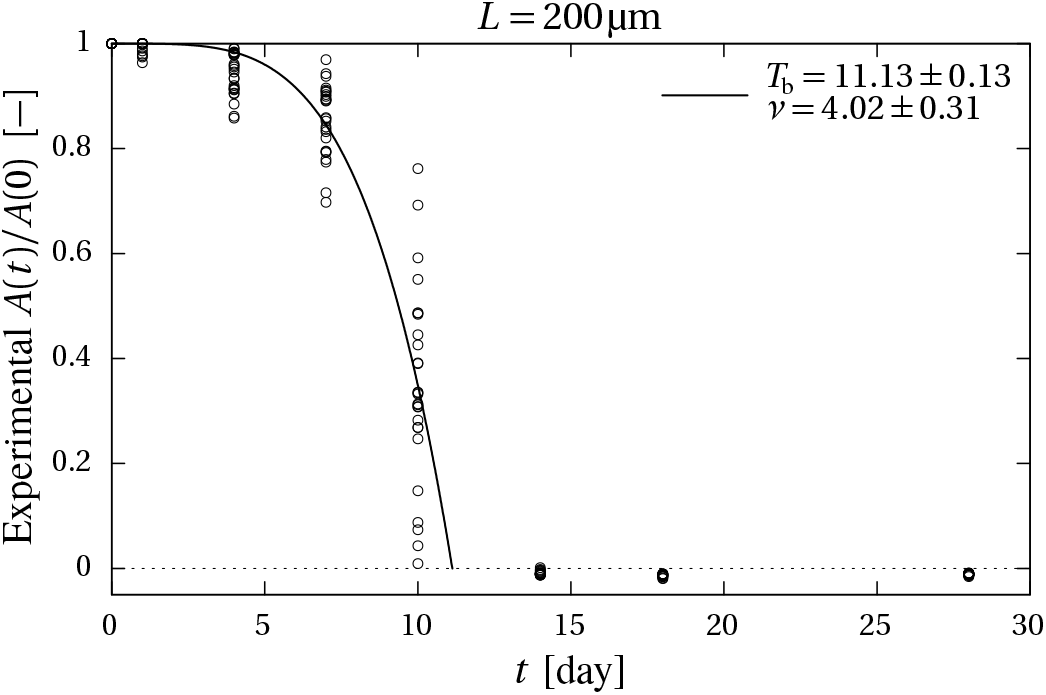
Experimental time course of pore area normalised by initial pore area (circles) and curve of best fit for Eq. (1) (solid line) for the mesh size *L* = 200 μm. The estimated time to bridge *T*_b_ ≈ 11.13 days is the time at which the fitted line intersects the horizontal axis (zero pore area). The parameters of best fit Tb and *v* are mentioned in the figure (± asymptotic standard error [57,56]). See supplementary figure S1 for the other mesh sizes.

### 2.2 Cellular reaction-diffusion model of tissue growth

Experimental images in Figure 2 suggest cell migration and cell proliferation are important mechanisms because we see clear evidence of the tissue forming by expanding into free space, and the density of cells behind the moving front increases with time. The experimental images show no obvious indication of cells changing shape or size during the experiments, and no indication of the presence of stressed or dead cells. Live/dead assays indicate that cell death is almost completely absent in these experiments (data not shown). Osteo-inducing factors in the cell culture media are excluded, so that cell differentiation is negated in the experiments. Since we are dealing with experiments involving large numbers of cells we work with a continuum reaction–diffusion model that describes cell density information rather than information about particular individual cells. Such reaction–diffusion models provide very good approximations of average behaviours even for low cell numbers [58]. Assuming that the tissue is composed of a uniform layer of cells in the out-of-plane (*z*) direction, and noting that the movement of the leading edge of the cell population during the bridging process maintains a well-defined sharp front (see Figure 1), we model the experiments using the Porous-Fisher equation,

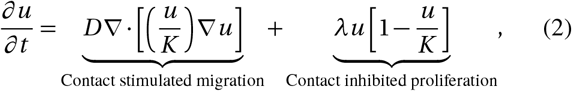

where *u*(*x, y, t*) ≽ 0 is the cell density (number of cells per unit area) that depends on time *t* and position (*x, y*) within the initial square pore; *D* > 0 is a cell diffusivity parameter; *λ*> 0 is a proliferation rate parameter; and *K* > 0 is the carrying capacity density. Both *D* and *λ* are intrinsic properties of the cells that represent the cells’ individual migratory and proliferative behaviour, respectively. Table 1 summarises the variables, parameters and their biological interpretation.

**Table 1.** Variables, parameters and interpretation in the Porous-Fisher model.

| Symbol | Description |
| --- | --- |
| $t$ | time [days] |
| $(x, y)$ | position in 2D Cartesian coordinate system |
| $u(x, y, t)$ | cell density [cell number per $\mu\text{m}^2$] |
| $T_b$ | time to bridge the pore [days] |
| $D$ | cell diffusivity [$\mu\text{m}^2/\text{day}$] |
| $\lambda$ | cell proliferation rate [/day] |
| $K$ | carrying capacity [cell number per unit area] |
| $L$ | initial pore size [$\mu\text{m}$] |
| $u_0$ | initial cell density at the pore boundary |
| $u_c$ | threshold density for tissue/pore boundary |
| $L_c$ | characteristic length scale $\sqrt{D/\lambda}$ |
| $\alpha^2$ | dimensionless parameter $D/(\lambda L^2) = (L_c/L)^2$ |

Equation (2) is the Porous-Fisher model that combines density-dependent diffusion to model contact-stimulated migration of cells, and logistic growth to model contact-inhibited proliferation of cells up to a carrying capacity density. These density dependences represent mechanistic crowding effects on collective cell proliferation and cell migration processes. The overall rates of these processes do not depend explicitly on pore size or pore shape, as cells are unlikely to sense geometrical features much larger than their individual size. However, pore size and pore shape constrain the availability of free space for cell motion and cell division, and these processes in turn influence cell density. We also note that the nonlinear diffusion term in Eq. (2) can be written as *D*(*u*) = *Du/K*, and since *D* (0) = 0, this leads to the formation of sharp fronts [59], as we see in the experimental images in Figure 1. Previous studies that have compared the performance of the Porous-Fisher model to the Fisher–Kolmogorov model have often observed that the Porous-Fisher model provides a better description of various types of *in vitro* experiments [46,52,48].

In the Porous-Fisher model the diffusivity *D* provides a measure of the motility of a cell within the experiment and the proliferation rate *λ* provides a measure of the rate at which a cell divides. Net cell proliferation is a decreasing function of density to reflect contact inhibition of proliferation. Where cell density is low, the per-capita proliferation rate is *λ*, and this per-capita proliferation rate decreases linearly to zero as the density approaches the carrying capacity *K*. The average effect of cell diffusion is to even out spatial heterogeneities of cell density. This model can be thought of as a simplification of a more realistic model of cell migration interacting with an extracellular matrix, see the supplementary information, Section S2. While *λ* can be relatively easily measured or inferred using estimates of the cell doubling time, the diffusivity is notoriously difficult to measure. Reported values of cell diffusivities in the literature vary over several orders of magnitude [60, 61, 42, 43, 62, 44].

Having now introduced the model and described its mechanistic processes and its cell behavioural properties, we can reformulate the key questions that we wish to explore in this study in this way:

i. Do estimates of *D* and *λ* vary across the cell bridging experiments performed with various pore sizes? If so, is there an optimal pore size that maximises tissue growth rate?
ii. Do estimates of *D* and *λ* vary in time within a single pore during pore closing? If so, can we relate these changes with changes in tissue geometry?

No variation of the rates *D* and *λ* across scaffolds of different pore sizes would indicate that cell-level behaviour is unaffected by the initial pore size and geometry. No variation of the rates *D* and *λ* in time would indicate that cell-level behaviour is unaffected by the current pore size and geometry. Without connecting a mechanistic model to the experimental data it would be very difficult to make such distinctions. It is important to note here that by disentangling the mechanistic behaviour and the cell-level behaviour of tissue growth in the experimental data, the mathematical model is then able to make predictions of tissue growth in other scaffold geometries that have not been realised experimentally. These predictions would allow the optimisation of scaffold topology based on numerical simulations, helping to narrow the space of experimental parameters to be tested experimentally. Taking such an *in silico* approach to experimental design and optimisation would lead to savings of time and experimental equipment.

#### 2.2.1 Nondimensionalisation

The cell proliferation and cell diffusion processes included in the mathematical model are described by three parameters: the diffusivity *D*, the proliferation rate λ, and the carrying capacity *K*. Two further constant parameters are associated with the initial conditions of the model: the initial density *u*_0_, and the initial pore size *L*. In addition, we introduce a threshold cell density *u*_c_ above which cells are considered part of the newly grown tissue, and under which they are considered part of the pore. That is, the tissue interface in the mathematical model is assumed to be given by the contour line *u*(*x, y, t*) = *u*_c_ where cell density equals *u*_c_. The three possible scalings that can be performed on the independent and dependent variables (space, time, and cell density) allow us to reduce the total number of independent parameters from five to two. Setting nondimensional variables of space, time, and density as 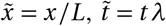, and 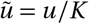, the governing evolution equation becomes

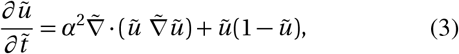

with the initial condition 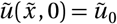, where

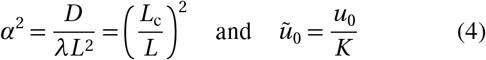

are the independent dimensionless parameters. These are supplemented by the dimensionless threshold density 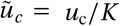 that identifies the tissue interface. The cellular processes of diffusion and proliferation define a characteristic length scale 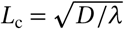 which corresponds to the average distance travelled by a cell by diffusion before it undergoes division. From this point on we work with dimensionless variables unless otherwise stated and we suppress the tilde notation. In essence, the nondimensional cell density is measured as a fraction of *K*, time is measured in units of λ^-1^, and space is measured in units of *L*. The combination of cell diffusivity and cell proliferation rate in *α*^2^ can be conveniently pictured as the ratio (*L*_c_/*L*)^2^.

In this work we interpret our experimental data using numerical solutions of the nondimensional Porous-Fisher model, Eq. (3). Numerical solutions of Eq. (3) are obtained using the method of lines and no-flux (symmetry) boundary condition on the unit square boundary. Full details of the numerical solution are outlined in the Supplementary Material. To connect the numerical solution of Eq. (3) with the experimental data on bridging time, we first use the numerical solution of *u*(*x, y, t*) to estimate the pore area as a function of time, *A*(*t*), by integrating numerically the region of the solution with *u*(*x, y, t*) < *u*_c_. We then define the time to bridge, *T*_b_, in the mathematical model as the time required for the normalised pore area *A*(*t*)/*A*(0) to fall below the tolerance *ε* = 1 · 10^-10^.

## 3 Results

Images in Figure 1 show snapshots of the cell bridging experiments, where we see that experiments with larger pore size *L* require a longer period of time to bridge than experiments with smaller pore size. For example, we see that of the nine pores shown with *L* = 200 μm, none are bridged at day 7, one is bridged at day 10, and all are bridged by day 14. Similarly, of the nine pores shown with *L* = 600 μm, none are bridged at day 14, one is bridged at day 18, and four are bridged at day 28. These results indicate that the time to bridge increases with *L*, but this does not indicate whether those cells in the experiments with larger *L* behave differently to those cells in the experiments with smaller *L*.

To provide insight into how the mechanisms in the experiments may vary with *L* we estimate the bridging time *T*_b_ from the experimental data and plot it as a function of *L* in Figure 3. We formalise the observed relationship between *T*_b_ and *L* by fitting a power law ∝ *L^μ^* to the data in Figure 3. The regression shows that bridging time is simply proportional to initial pore size, since we find that *μ* ≈ 0.999094, which is remarkably close to unity. To quantify the linearity of the relationship between bridging time and initial pore size, we also fit the data using a linear regression in Figure 3. The data clearly suggests a constant, linear relationship between bridging time *T*_b_ and inital pore size *L* for all *L* considered, which we write as

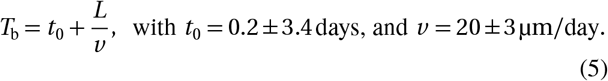

**Figure 3.**
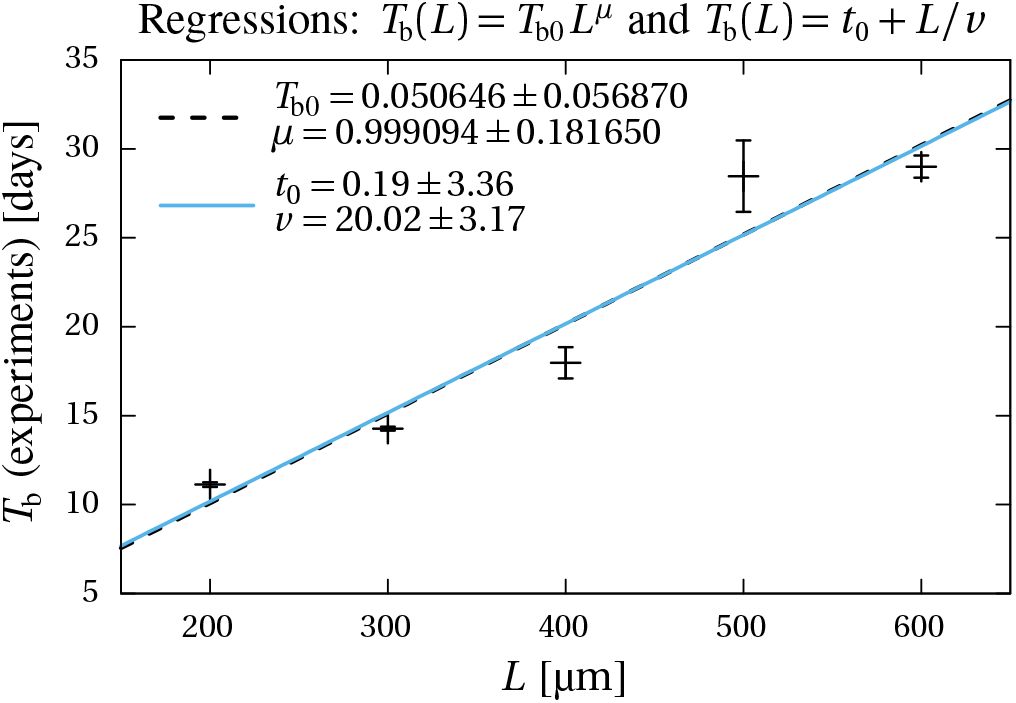
(Color online.) Experimental bridging times vs pore size, estimated from Figure 2 and Supplementary Figure S1 (pluses). Error bars corresponds to the asymptotic standard error of the determination of *T*_b_ (see text). These experimental data are fitted with a power-law regression (interrupted black line) and a linear regression (solid blue line). The coefficient of determination quantifying the degree of linear correlation between *T*_b_ and *L* for the linear regression is *R*^2^ ≈ 0.93.

The intercept *t*_0_ corresponds to the time taken to bridge a pore of zero size (*L* = 0). Therefore, the estimate of *t*_0_ ≈ 0 is expected. The linear regression parameter *v* can be interpreted as an average velocity of tissue progression, and the estimate of *v* ≈ 20 μm/day is consistent with the time and length scales in the experimental images shown in Figure 1. We will now investigate the implication of the relationship (5) on the potential values of the parameters *D* and *λ* of the model.

To see how the mathematical model relates to the experimental observations in Figure 1 we present numerical solutions of Eq. (3) on the unit square in Figure 4. The still frames in Figure 4a show the evolution of the system with various values of *α*^2^ = 0.01,0.02 and 0.05. The evolution of the numerical solutions is consistent with the experimental observations in Figure 1 since we see that during the early part of the experiment, the new tissue is formed and the shape of the infilling tissue is relatively square. As the experiment proceeds, the shape of the infilling tissue becomes increasingly rounded before closing at some point. In Figure 4, contour lines of cell density are shown every 0.1 increment. The red contour shows the interface that we take to represent the boundary of the inward-growing tissue, i.e., the contour line at density *u_c_* = 0.5. Comparing the evolution of this contour for the different parameter values in Figure 4a shows that the bridging time is a decreasing function of *α*^2^ = (*L*_c_/*L*)^2^. In other words, the greater the distance *L*_c_ travelled by cells before they undergo division, the lesser the bridging time. Analogous results in Figure 4b show solutions for various values of *u*_0_ = 0.8, 0.5 and 0.2. Again, the features of the evolution of the numerical solutions are consistent with the experimental images in Figure 1 and here we see that the bridging time is a decreasing function of *u*_0_ at constant α, i.e., the more cells per unit surface we start with initially in a pore, the lesser the bridging time, which is intuitively reasonable. It is clear by inspection of the contour lines in Figure 4 that similar conclusions hold for other choices of *u_c_*.

**Figure 4.**
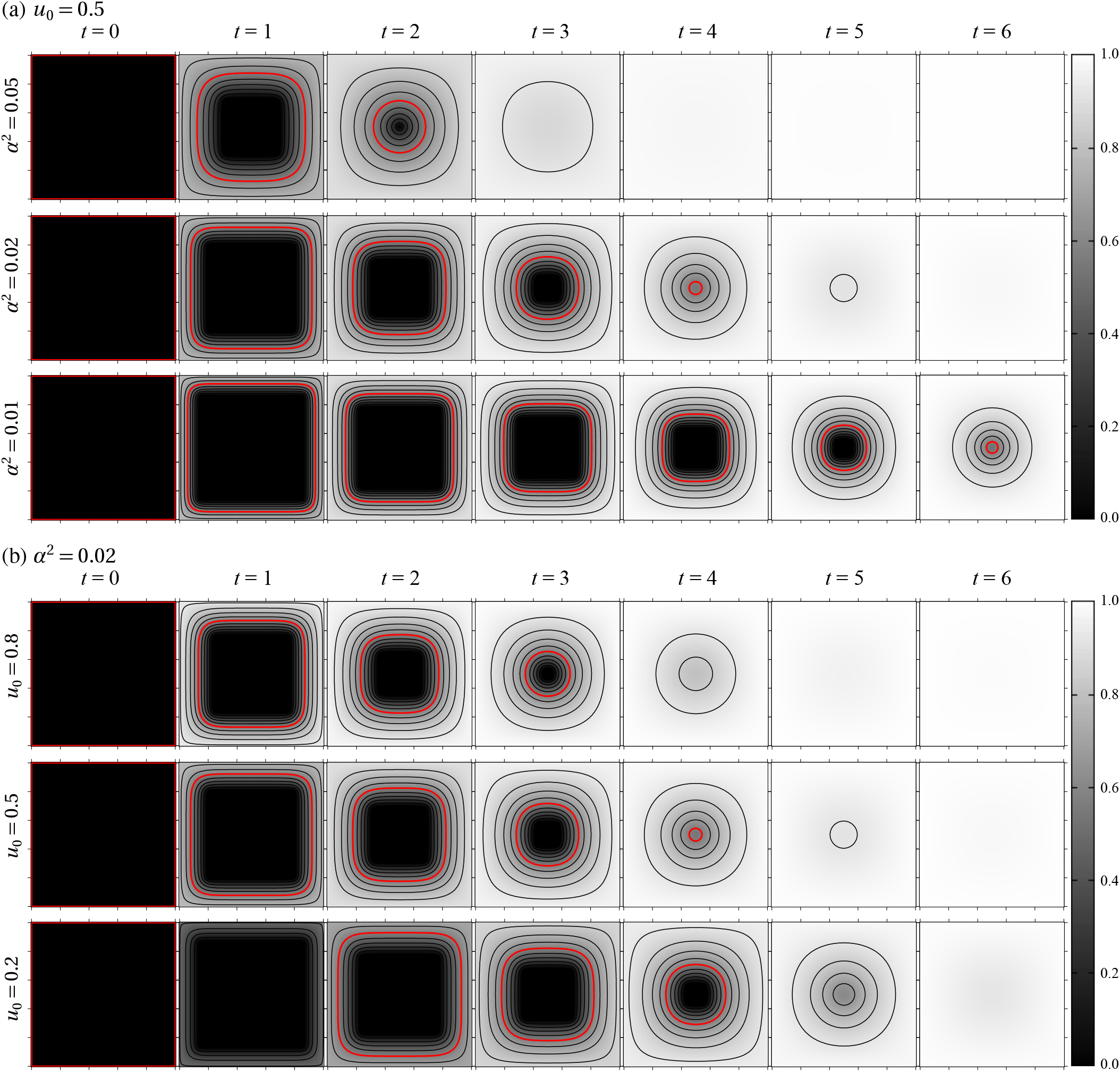
(Color online.) Still frames from the nondimensionalised mathematical model at even time intervals for different combinations of the parameters *α*^2^ = (*L*_c_/*L*)^2^ and *u*_0_ (rows). (a) The initial density is set to *u*_0_ = 0.5 and *α*^2^ varies; (b) The parameter *α*^2^ is set to *α*^2^ = 0.02 and the initial density *u*_0_ varies. The threshold density (red) is set to *u*_c_ = 0.5. Isolines of constant (nondimensional) density are shown every *Δu* = 0.1. The middle rows in (a) and (b) are identical and correspond to *α*^2^ = 0.02, *u*_0_ = 0.5.

A closed-form solution to Eq. (3) is not known for this particular geometry, so we use numerical estimates to describe Tb as a function of initial pore size *L*. Using the dimensionless model we plot *T*_b_ as a function of *L* in Figure 5a. Consistent with the experimental data, we observe that *T*_b_ is a linear function of *L* when *D* and *λ* are maintained at fixed values:

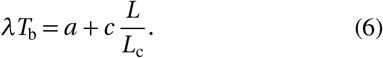

**Figure 5.**
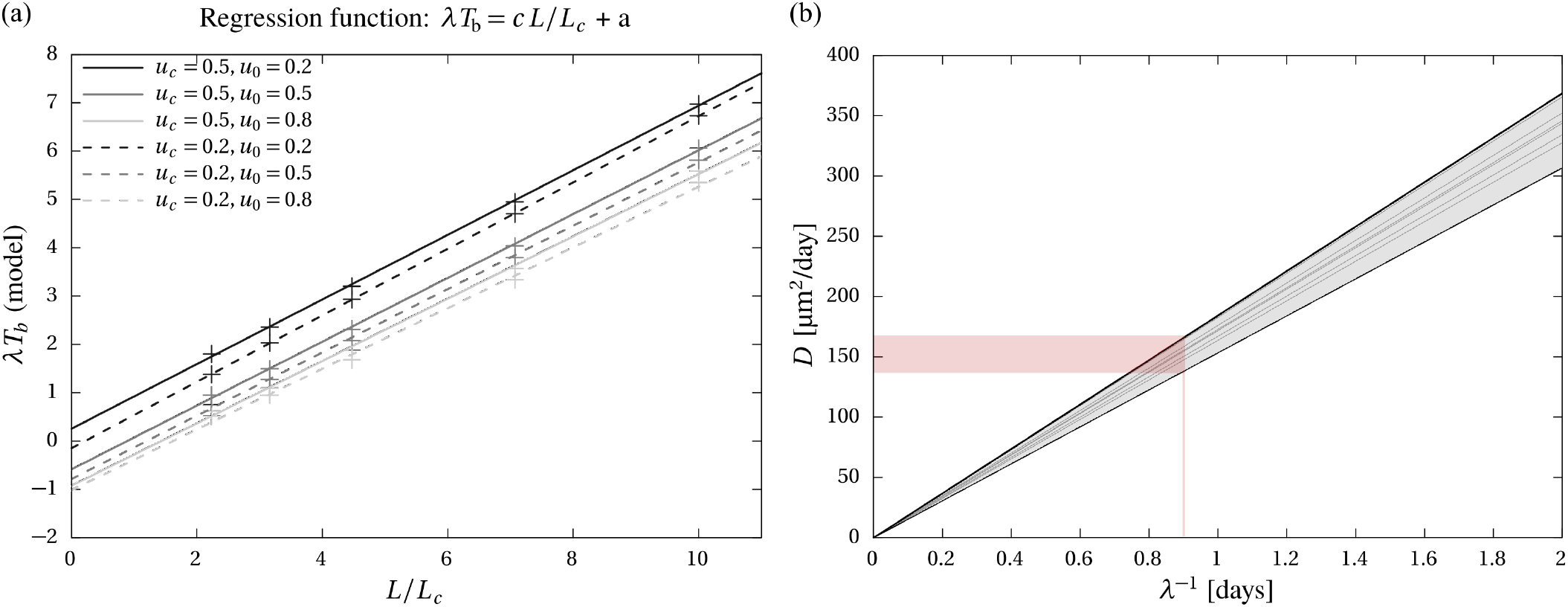
(Color online.) (a) Bridging times vs pore size in the mathematical model. Individual simulation results are shown as pluses (+) for a number of choices of *u*_c_ and *u*_0_. Corresponding lines of best fit satisfying Eq. (6) are shown as solid lines (*u*_c_ = 0.5) and interrupted lines (*u*_c_ = 0.2). Regression parameters are listed in Supplementary Table S4. In this plot, time is measured in units of λ^-1^ and pore size (∝ 1/*α*) is measured in units of *L*_c_. (b) Diffusivity *D* vs cell division period λ^-1^ compatible with the experimental data. The slopes *D*λ of the grey lines are mentioned in Supplementary table S4; they range from 153 μm^2^/day^2^ to 184 μm^2^/day^2^ (grey shaded area between the solid black lines). The red shaded area displays the range of diffusivities compatible with λ= 1.1/day.

This linear relationship holds for any value of the carrying capacity *K*, the initial condition *u*_0_, and the threshold *u*_c_, but the slope *c* and intercept *a* vary with *u*_0_ and *u*_c_ (Figure 5a, regression lines; Supplementary Table S4). We find that the slopes *c* of the regression lines in Figure 5 are very consistent, varying between 0.63-0.69, while the intercepts *a* vary between −1.02 and 1.18 and depend on the initial density. Starting the simulations with a different initial density affects the time to bridge by a constant offset which is the same for all initial pore sizes. To connect these results with the experimental relationship between bridging time and initial pore size in Eq. (5), we rewrite Eq. (6) in dimensional form as

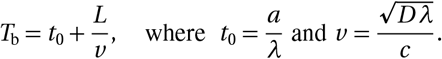

In particular we obtain

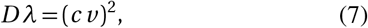

which provides a functional relationship between values of the cell’s intrinsic properties *D* and *λ* that are consistent with a linear dependence between bridging time and initial pore size in the model. We plot the relationship (7) between *D* and *λ* in Figure 5b by combining the regression estimate of *v* from the experimental data in Eq. (5), and the regression estimates of *c* from the numerical simulations with the set of values *u*_0_ and *u*_c_ listed in Supplementary Table S4. Figure 5b shows that on average, the model matches the experimental data for all pore sizes whenever the intrinsic cell diffusive behaviour and cell migratory behaviours represented by *D* and *λ* are (i) constant, i.e., independent of time and independent of initial pore size; and (ii) such that *Dλ* ≈ 170 μm^2^/day^2^. Given an estimate of *λ* we can determine an estimate of *D*. For example, the doubling time of MC3T3-E1 osteoblast cells are approximately 15h [55], giving *λ* ≈ 24log(2)/15 ≈ 1.1/day. The relationship in Figure 5 indicates that this corresponds to *D* ≈ 140-170 μm^2^/day. This result is remarkable in its consistency. For example, it is worth noting that estimates of *D* in the literature vary over two orders of magnitude (see [44, and refs cited therein]), so the fact that we obtain such tight estimates here points to two important results: (i) our experimental protocols are remarkably consistent; and (ii) the experimental observations are extremely well described by the Porous-Fisher model with a constant set of parameters.

## 4 Discussion

There is experimental evidence that the mechanical stress experienced by a cell can affect its behaviour, such as its propensity to undergo cell division [12, 24, 25]. Previous experiments growing epithelial cell sheets suggest that the shape of the tissue regulates patterns of proliferation, and that this regulation may be induced by local mechanical stress [12]. In engineered tissue scaffolds, tissue grown by osteoblast-derived cells in pores of different shapes is also observed to be regulated by geometry. The local rate of growth is found to correlate with the curvature of the tissue [13–15], and thought to be due to tissue surface tension driving cell proliferation [14–17]. Phenomenological models that describe these scaffold experiments assume that the evolution of the tissue interface is governed by mean curvature flows, by analogy with surface-tension-induced mean curvature flows that arises in the evolution of bubbles in fluid mechanics. In these phenomenological models of tissue growth, cells are not represented explicitly, and neither is the volumetric growth of the tissue. Only the interface of the tissue is evolved, with no consideration of the correlation between volumetric tissue growth, and cell proliferation or tissue synthesis.

In our cell bridging experiments in Figure 1, the rounding off of the tissue interface at corners of the initial pores suggests that tissue grows preferentially where curvature is high, consistently with other studies [13–15, 17]. Because the tissue interface evolves through a series of shapes with increasing curvature, it is not obvious why the time required to bridge the pores in these scaffolds varies linearly with initial pore size (Figure 2). It could be expected that cell-level proliferation would vary not only in time, but also depending on the size *L* of the initial pore, since the overall scale of a pore affects its average curvature.

The mathematical model of cell diffusion and proliferation that we propose enables us to model volumetric tissue growth from cellular mechanisms. This allows us to analyse our experimental observations in terms of the cell diffusive behaviour (encapsulated in the cell diffusivity *D*) and the cell proliferative behaviour (encapsulated in the cell proliferation rate *λ*). As seen in Figure 4, the model agrees very well qualitatively with the different shapes taken by the tissue during its growth. New tissue forms at an increased rate near corners of the initial pore, so that the interface gradually takes on a more circular shape until bridging occurs. Importantly, the time required to bridge the pores in the mathematical model depends linearly on initial pore size *L* (Figure 5a) when it is assumed that cell diffusivity *D* and cell proliferation rate *λ* (i) are constant throughout the simulation (independent of time); and (ii) take on the same values across pores of different initial size. These results suggest that the mechanistic crowding effects represented by the density dependence of collective cell proliferation and cell diffusion in Equation (2) capture the dynamics of tissue growth in different pore geometries very well. Neither the initial pore size nor the transient pore size or geometry affects the individual cell proliferative behaviour (corresponding to *λ*) and individual cell diffusive behaviour (corresponding to *D*) in these scaffold experiments. The mechanistic processes of proliferation and diffusion of the model considered in Eq. (2), i.e., contact-stimulated migration and contact-inhibited proliferation, are sufficient to describe the evolution of the tissue without requiring additional dependences upon geometrical features of the pores.

Despite the independence of the cell’s proliferative and diffusive behaviour upon local curvature in the model, the produced tissue exhibits curvature-dependent growth rates. An often overlooked factor of the geometric control of tissue growth is the mechanistic influence of tissue crowding or spreading in confined spaces. The progression rate of the tissue interface is determined both by how much new tissue is produced locally, and by the availability of space around where this tissue is produced [36, 27, 37]. The former is related to cell behaviour, but the latter is a purely geometric influence. Space in a concave region of the tissue quickly runs out, and so to accommodate new tissue material, the tissue invades more space in the normal direction than in the lateral directions, leading to faster progression rates. In contrast, there is more space available to accommodate new tissue in a convex region, so that tissue progression in the normal direction is slower as the new tissue needs to fill space laterally too. When new tissue production is confined in a very narrow band near the tissue interface (a situation called ‘surface growth’, such as that which occurs during bone formation, and the growth of seashells [63]), the mechanistic influence of space constraints on growth rates lead mathematically to a type of hyperbolic curvature flow in which the normal acceleration of the tissue is proportional to curvature [36, 27, 37]. In the Porous-Fisher model considered here, proliferation is not strictly confined to the tissue interface. However, the logistic, contact-inhibited proliferation term in Eq. (2) means that in the pore region, where cell density is zero, and in the tissue region, where cell density approaches carrying capacity, there is no cell proliferation, and therefore no tissue growth locally. The local production rate of the tissue induced by the logistic proliferation term is highest where the normalised cell density is 0.5, corresponding to *u_c_* and to the red contour in Figure 4. Geometric control of tissue growth only occurs where there is differential growth rate, i.e., where the progression rate of the tissue interface is different in one region compared to another. Clearly, differential growth rates in the Porous-Fisher model arise where there are strong spatial heterogeneities in cell density, which is precisely in a relatively narrow band around the tissue interface marked in red in Figure 4. While cell proliferation also occurs away from the tissue interface, or after bridging has occured (e.g. top row in Figures 4a,b), there is little influence of geometry on tissue growth in these regions because density heterogeneities are small. In the experiments, increases in density occur after bridging as well, until reaching a maximum. For example, at Day 18 with *L* = 400 μm in Figure 1, some scaffold pores retain the memory of a circular closing pore by having lower density there.

We note here that while the Porous-Fisher model has sharp fronted solutions, the definition of a continuum density relies on the definition of a local averaging window. A nonzero density of cells *u* outside the tissue region corresponds to taking parts of the cellular tissue into this averaging window. Here we chose to emphasise the contour *u*_c_ = 0.5 as the tissue interface since this is midway between the minimum and maximum densities in the nondimensionalised model. Other numerical choices of *u*_c_ are possible. Figure 5a shows that the main result of the paper, i.e., that the time to bridge is a linear function of pore size, holds regardless of the choice of *u*_c_.

The mathematical model presented is deterministic. As such, it does not account for the variability seen in the experimental images and data in Figures 1–3. Model results should be interpreted in an average sense. From our experimental observations, the largest factor of pore bridging variability comes from whether, and when, the process of pore closing is initiated. This often depends on the location of specific pores within a scaffold, which may be due to experimental confounding factors, e.g., related to cell seeding and nutrients, and culture adaptation. Once pore closing is initiated, our experimental observations suggest that the evolution of the tissue in the pores follows a more deterministic course. Clearly, the time at which pore closing is initiated influences the time to bridge directly. Our mathematical model assumes pore closing to be initiated immediately. However, low initial densities *u*_0_ in the model act mostly to delay the time to bridge by a constant period of time (Figure 5a), and may thus be considered a good proxy to model experimental delays in pore closing initiation.

The nondimensionalisation of the model shows that, up to factors scaling physical units, only two parameters are independent: the initial density *u*_0_ and the parameter *α* = *L*_c_/*L*, which combines cell-specific parameters related to proliferation and diffusion, and initial pore size. The initial density mostly shifts time to bridge *T*_b_ by a constant offset independent of initial pore size (Figure 5a). The average distance *L*_c_ travelled by a cell before it undergoes division measures a trade-off between cell diffusion and cell proliferation for the behaviour of a single cell. Different choices of cell proliferation and cell diffusivities that combine into a same value of *L*_c_ will generate exactly the same succession of cell density profiles and tissue interfaces in space, except that these will be reached at different times due to the scaling of time units by λ^-1^. Scaling analysis alone does not predict that time to bridge *T*_b_ should be proportional to initial pore size. This result is obtained by simulating the model with pore sizes ranging from about two to ten times the characteristic length scale *L*_c_ (Figure 5a). The linear relationship between *T*_b_ and *L* in the model shows that overall, tissue invasion of a pore space of linear size *L* occurs with an average speed 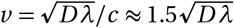. The linear relationship between bridging time and initial pore size in the experiments suggests that this average speed is approximately 20 μm/day for any initial pore size, see Eq. (5). This result indicates that the dynamics of tissue invasion in these pores is mostly geometric; there is no optimal pore size at which the rate of tissue progression into the pore is highest. Time to bridge is simply proportional to the geometric size of the pore. This is consistent with our conclusion that cell behaviours do not seem to be affected by the initial pore size significantly.

Several combinations of diffusivity *D* and cell division period λ^-1^ are compatible with the experimental data. The slopes of the curves *D* vs λ^-1^ in Figure 5b are remarkably consistent across the numerical simulations, which is due to the consistency in slope of the linear relationship between bridging time and initial pore size. Cell doubling times for the cell line employed in the experiments are known to be very consistent [55]. There is more experimental uncertainty about cell diffusivity [60, 61, 42, 43, 62, 44]. Figure 5b integrates information both from the experimental data and from the mathematical model, and enables us to estimate the range of diffusivities for this cell line in the pore bridging experiments conducted in Figure 1. With the estimated parameter values *λ* ≈ 1.1/day and *D* ≈ 140-170 μm^2^/day, the estimated average distance travelled by a cell in the bridging experiments before it undergoes division is *L*_c_ ≈ 12 μm. This is consistent with our observation that the cells remain closely bound with each other throughout the experiments [53]. We note that these estimates of cell diffusivity are similar to the estimates 50–130 μm^2^/day found for prostate cancer cells in scratch assays [44]. These estimates are remarkably consistent given that estimates of *D* typically vary over several orders of magnitude in the literature [44].

Porous-Fisher models are known to lead to travelling waves for certain initial conditions, boundary conditions and geometries [38, 42, 43]. In one spatial dimension, stable travelling waves progress at speed 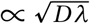. This suggests that the linear dependence of time to bridge *T*_b_ with initial pore size *L* is an approximation due to the fact that the tissue interface propagates into the pore as a travelling wavefront at constant speed for most of its trajectory, except at the onset of pore closing where the wave requires sufficient cell density buildup to establish itself [64]. This observation suggests that other pore shapes with linear size *L* ≿ *L*_c_ would also bridge with a time that increases linearly with *L*. Figure 6a shows a simulation of the Porous-Fisher model of tissue growth in a rectangular pore shape of size ratio 2:1. Figure 6b confirms that time to bridge increases linearly with initial pore size also for rectangular pores. The slope *c* of the linear regression is similar to the slopes *c* reported in Figure 5a, meaning that on average, the tissue front propagates at a similar velocity of approximately 20 μm/day.

**Figure 6.**
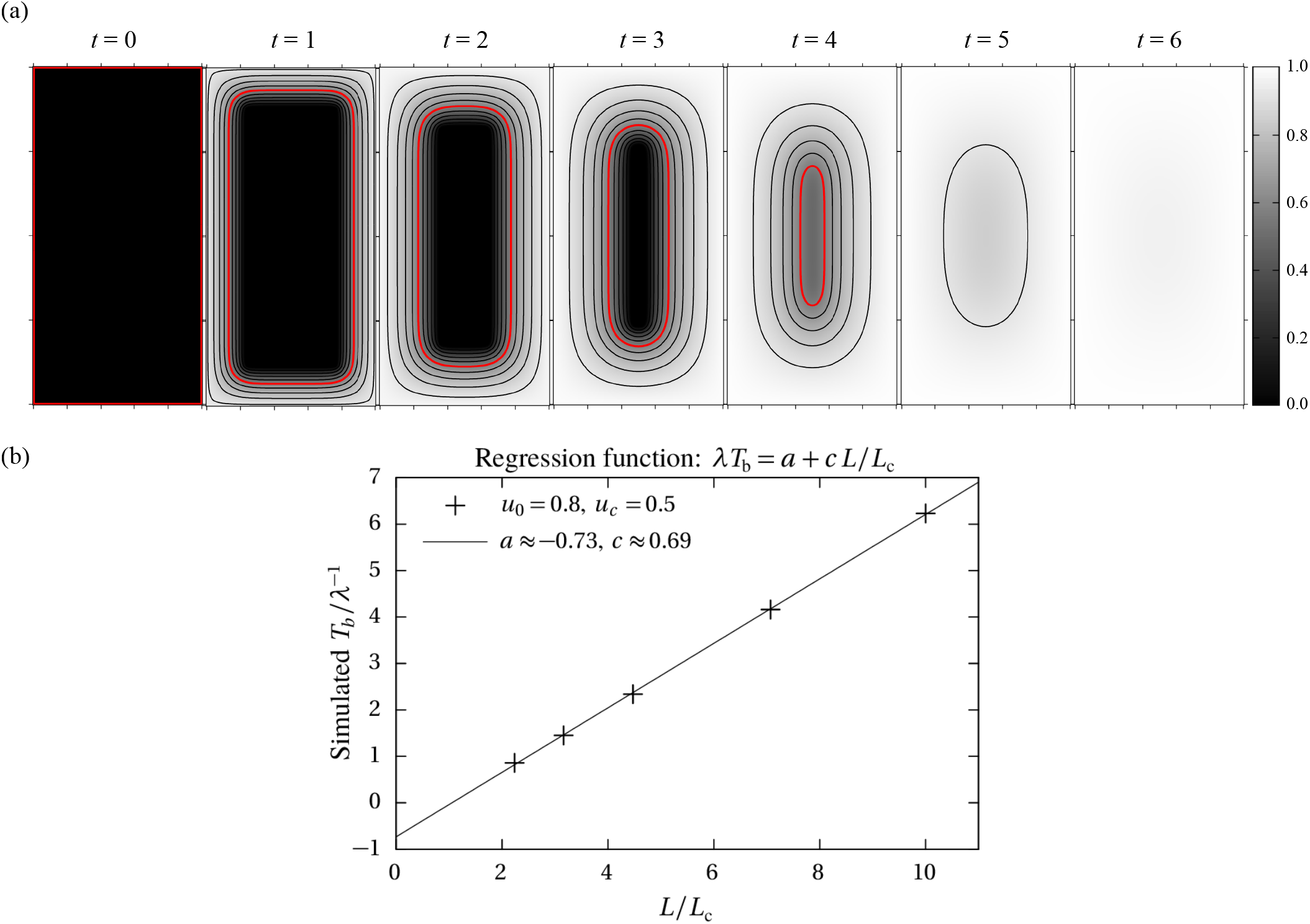
(Color online.) Porous-Fisher simulations on a rectangular initial pore. (a) Still frames from the nondimensional model at even time intervals for *α*^2^ = 0.02, *u*_0_ = 0.8, and *u*_c_ = 0.5; (b) Time to bridge for rectangular pore of different sizes: simulation data (crosses) with linear regression line (solid line).

The key result of our experimental measurements and numerical modelling is that *T*_b_ increases linearly with *L* is limited to some intermediate range of *L*. For example, if our experiments were repeated with *L* sufficiently small that *T*_b_ was much less than the timescale of proliferation would mean that cell migration alone would control *T*_b_ rather than combined cell migration and cell proliferation. In this limit we would expect *T*_b_ would be proportional to *L*^2^ rather than *L* [65]. Similarly, if we were dealing with an extremely large pore size *L*, or less robust or multipotent primary cells or cell lines, then some other mechanisms such as cell death or cell differentiation could play an important role and so *T*b may not scale linearly with *L* in this case. Rather, the conclusions of our investigation are relevant for pores with an intermediate length scale, 200 < *L* < 600 μm and some care ought to be exercised when extrapolating our results to pores that are either sufficiently small or sufficiently large, or that use different cell types.

## 5 Conclusion and Future Work

In this paper, we propose a simple reaction–diffusion model of cells to analyse in detail the experimental observation that the time to bridge pores in 3D printed scaffolds increases linearly with initial pore size. The mathematical model includes two cellular mechanisms at play during tissue growth: cell proliferation, and cell transport. The model contrasts with previous phenomenological models of the geometric control of tissue growth based on mean curvature flow, which do not consider cells explicitly and which assume an explicit dependence on curvature.

Our mathematical analysis suggests that the linear dependence observed experimentally between time to bridge and initial pore size is solely due to geometric crowding effects and the spatial scale of the pore space in which the tissue grows. The cell behaviours, encapsulated in the model by the diffusivity *D* and the cell division rate *λ*, do not need to be adjusted for the model to recover a linearly increasing bridging time with pore size. Furthermore, it is anticipated from the model that pores of different shapes may also bridge in a time that scales linearly with initial pore size. These results suggest that there is not an optimal pore size that maximises the rate of tissue progression within reasonable size limits leading to tissue bridging.

While the Porous-Fisher model allows proliferation to occur wherever cells occupy space with lower density than the carrying capacity, in effect, geometric control of cell proliferation is concentrated in a region near the tissue interface where there are large density inhomogeneities. The Porous-Fisher model is similar in this respect to the cellular models of surface growth of Refs [36, 27, 37]. A common feature in these cell-based models is that an influence of local curvature on tissue growth rate is not necessarily indicative of changes in cell-level behaviour. To determine cell-level behaviour, it is important that experimental data is examined with the help of mathematical models to factor out the mechanistic influences of space constraints induced by tissue crowding [27], as we have done here to conclude that cell-level proliferation and diffusivity parameters are independent of pore size.

Our results suggest that the dynamics of tissue growth by the MC3T3 cell line in 3D printed scaffold pores is dominated by mechanistic proliferation and diffusion mechanisms that rely only on local sensing of cell crowding, and not on local sensing of geometric features of the tissue that may span many cell diameters. While there seems to be no or little cell-level geometric regulation in our experiments, there is emergence of tissue-level geometric regulation arising from cell crowding effects in confined spaces. The elucidation of the mechanistic and cell-level diffusive and proliferative mechanisms required to match our experimental data is an important outcome of our study. It allows the prediction of the evolution of MC3T3-cell-produced tissue in scaffold pores of new geometries, and can thus help design optimal scaffold pore shapes to meet conflicting constraints in both space and time, such as the requirement to bridge pores quickly while maintaining some degree of permeability [18].

A faithful mimicry of tissue or implant growth in biologically more complex environments would involve a consideration of material properties, three dimensions, and several tissue types with paracrine interactions. Our study focused on the growth of tissue to bridge scaffold pores in a highly reproducible setup that does not include more complex features relevant to holistic tissue models, but that is particularly prone to mathematical analysis. Our use of thin bilayer scaffolds and frequent medium feeding circumvents issues of nutrient and metabolite exchange, as well as issues of three-dimensional tissue growth. Our use of a single, robust, proliferative MC3T3 cell line avoids patient-to-patient variability, loss of proliferative capacity, and multicellular interactions. The mathematical model does not capture some more complex cellular behaviours, including an initial culture lag phase or differentiation. For example, it is experimentally observed that very large pores are unlikely to bridge at all, whereas the Porous-Fisher model presented in this paper will bridge pores of any size. The diameter of the scaffold fibers could also play a role for the spatio-temporal organisation of the tissue created [20], particularly in the early stages of pore infilling. More detailed mathematical models with further comparisons to experimental data are needed to alleviate these limitations and will be the subject of future works.

The model presented here is simple in both the experimental data required and prediction provided. The remarkable linearity between pore size and the time to bridge, coupled with the predictive accuracy of this reaction-diffusion model, posits that complex relationships between single cell behaviour or substrate curvature may be unnecessary to identify useful tissue engineering design equations for the additive manufacturing era.

There are many interesting extensions of the work presented here, both experimentally and theoretically. The first extension would be to consider very similar experiments on different shaped pores, such as the rectangular pore that we studied theoretically in Figure 6. Extending our experimental design to include rectangular, circular, triangular and cross-shaped (nonconvex) pores is of interest and some of our previous *in vitro* work using much simpler two-dimensional wound healing assays [46, 47] have begun to explore these details. The investigation of nonconvex shapes is of particular interest as previous studies have shown that the average rate of tissue growth in porous scaffolds depends on whether pores are convex or nonconvex [13, 21].

More direct experimental measurements of single-cell proliferative and migratory behaviours in their population environment at different locations and time points during the pore bridging experiments would help validate the importance of cell crowding effects for the geometric control of tissue growth. These kinds of experimental data would also be useful to determine whether generalisations of the Porous-Fisher model with time-dependent coefficients are required [61, 67], such as a time dependent diffusivity *D* (*t*) and a time dependent proliferation rate, *λ*(*t*), such that the product *D*(*t*)*λ*(*t*) is maintained constant. We have not considered this possibility here because there are no obvious trends in our experimental observation that motivate this kind of extension, however, this is a potential avenue for future consideration.

Another interesting extension would be to extend the mathematical modelling of the cell density presented here using a full three-dimensional analogue of the Porous-Fisher model. This would involve working with a three-dimensional analogue of Equation 2 for cell density, *u*(*x, y, z, t*). This approach would be far more challenging because we would need to specify much more information that is currently uncertain. For example, this approach would require us to specify the initial condition, *u*(*x, y, z*,0), as a function of vertical position as well as specifying boundary conditions on the lower and upper surfaces of the domain. Both of these details are presently uncertain and so this partly justifies working with a simpler two-dimensional vertically averaged model which is known to be accurate under the conditions we consider here, namely that the vertical length-scale is much smaller than the horizontal length scale [66]. Nonetheless, despite the challenges and uncertainties of working with a full three-dimensional model, it would be interesting to systematically compare the performance of a full three-dimensional modelling approach with the current two-dimensional modelling approach.

Finally, repeating our experiments and modelling work with different cell types is also of high interest, particularly where combinations of cell types involve paracrine interactions, since this situation represents more closely tissue or implant growth in biologically more complex environments.

Key algorithms used to generate numerical simulations of the mathematical model are available on GitHub.

## 6 Acknowledgements

This research was supported by the Australian Research Council (DP180101797, DP200100177) and the Centre for Biomedical Technologies, Queensland University of Technology (QUT). MCA acknowledges support from the Queensland Government through an Advance Queensland Industry Research Fellowship. We appreciate the helpful comments from the three anonymous referees.

## S1 Experimental bridging times

The time to bridge for each pore size is estimated from experimental data on pore coverage area versus time [53]. Figure S1 shows plots of pore area versus time normalised by initial pore area. Each data point comes from the evaluation of a single pore in an experimental scaffold. Nine pores are evaluated in each scaffold, but the data at different time points comes from different scaffolds [53]. At each time, there is a total of 9 × 3 = 27 data points for each mesh size *L*, except at *t* = 18days were there are 9 × 2 = 18 data points for the mesh sizes *L* = 200,400,500 μm. All the data from time points prior to bridging are used to fit curves 1 -(*T*_b_/*T*_b0_)^*ν*^ for each mesh size *L* using nonlinear least squares. (Bridging is assumed to have occurred when normalised pore area data is below a tolerance of 0.02.) The fitted parameter *T*_b0_ provides an estimate of the time to bridge. Estimates of the standard error on this fitted parameter are given by the asymptotic standard error calculated during the fitting procedure [57, 56].

**Figure S1.**
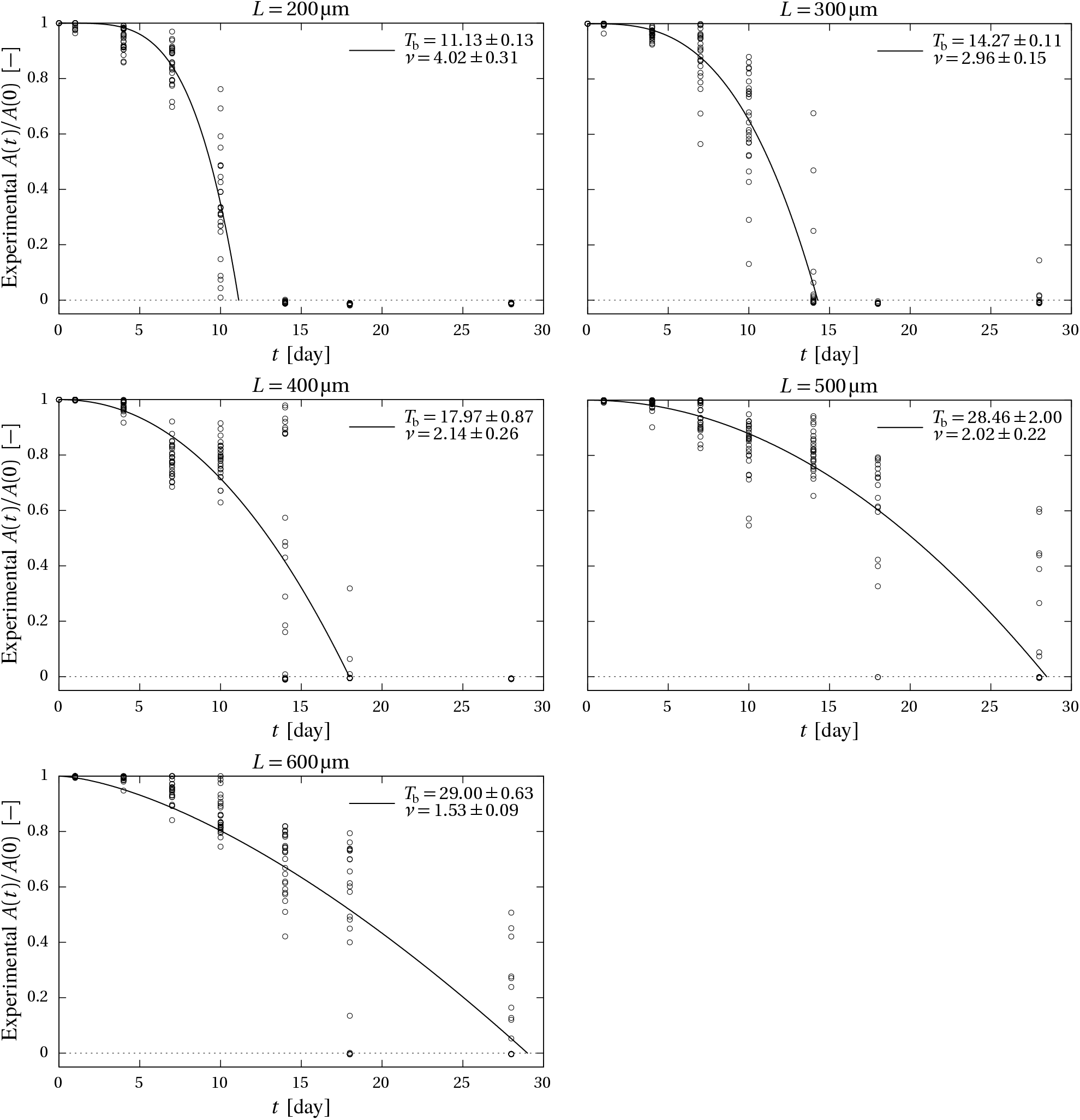
Estimation of time to bridge in the experiments. These figures show plots of the experimental pore area versus time, normalised by initial pore area (circles), as well as curves (solid lines) fitted on all the data points prior to bridging. Fitted parameters *T*_b_ and *v* and their asymptotic standard error [57, 56] are mentioned in the figure legends.

## S2 Mathematical model formulation

In the main document we comment that the Porous-Fisher model can be thought of as a simplification of a more realistic model where cells migrate in response to a substrate, such as the extracellular matrix (ECM). In this case we can think of cells producing ECM, which then facilitates the migration of those cells, a mathematical model of this kind of interaction between a population of cells, with density *u*(*x, y, t*), and ECM with density *e*(*x, y, t*), can be written as:

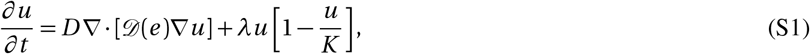

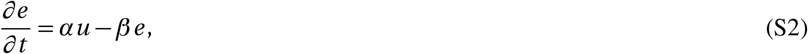

where *D* > 0 is the cell diffusivity, *λ* > 0 is the cell proliferation rate, *K* > 0 is the carrying capacity density, *α* > 0 is the production rate of ECM and *β* > 0 is the decay rate of ECM. In Equations S1–S2 we suppose that the diffusion of cells is coupled to the presence of the ECM through the nondimensional nonlinear diffusivity function, 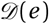, which we have not specified, but it would be reasonable to assume that 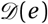 is an increasing function of the ECM, 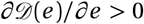. A simple and instructive choice would be 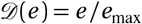, where *e*_max_ is some maximum characteristic density of ECM. Setting 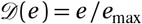 indicates that there is no migration in the absence of ECM and an increasing density of ECM enhances migration. To connect this model, Equations S1–S2, with experimental data we would ideally have observations of both the cell population, *u*(*x, y, t*), and the density of ECM, *e*(*x, y, t*). Since we do not have this kind of data in our experiments we invoke a standard simplification by assuming the time scale of ECM production and decay is faster than the time scale of cell migration and cell proliferation. This kind of quasi-steady assumption is widely invoked whereby the dynamics of some kind of signalling molecule is thought to reach equilibrium conditions faster than some population of cells [69, 71].

To simplify Equations S1–S2 we set *∂e/∂t* = 0, giving *u* ∝ *e*, so that we may write 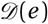 as 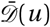, where 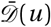 is an increasing function of *u* such that 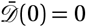, with an obvious candidate being 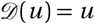, thereby recovering the Porous-Fisher model, which we write as

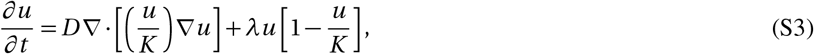

where we have written 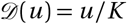 so that the nonlinear diffusivity function is non-dimensional. In this section we provide a very brief justification of the use of the Porous-Fisher model in terms of simplifying a multi-species model of cell migration and cell proliferation with a quasi-steady assumption simplifying the way that cell migration is coupled to ECM. This kind of assumption is one way to motivate the use of the Porous-Fisher model. Other arguments have been used to support the use of the Porous–Fisher model include arguments based on: (i) heuristic reasoning about the role of crowding-induced directed motion [49]; (ii) effects of cell-to-cell crowding and cell shape [51, 68]; (iii) or geometric arguments relating to asymptotic observations of hole-closing near to the time of closure [48]; and (iv) heuristic arguments about the observation of sharp fronts in experimental images [42, 43].

## S3 Numerical solution

Numerical solutions of the Porous-Fisher model are obtained by writing Equation S3 as

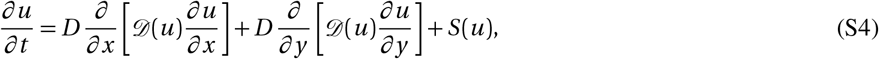

where we have written the model in terms of a general nonlinear diffusivity function, 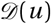 and a general source term, *S*(*u*). For our purposes we set 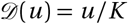 and *S*(*u*) = *λu*(1 — *u/K*). Our aim is to obtain numerical solutions of Equation S4 on the square domain *Ω* = {(*x, y*),0 < *x* < *L*,0 < *y* < *L*}. For convenience we assume that the origin is at the lower left corner of the domain and we discretise *Ω* on a spatially uniform finite difference mesh with mesh spacing *h* > 0. We index the mesh in the usual way so that the coordinates of each mesh point are (*x_i_, y_j_*, with *i* = 0,1,2,…, *I* and *j* = 0,1,2,…, *J*. Since we always consider square domains we have *I = J* = *L*/Δ, giving a finite difference mesh with a total of (*I* +1)^2^ nodes.

We solve Equation S4 using a standard method of lines approach so that at each internal mesh point we approximate Equation S4 by

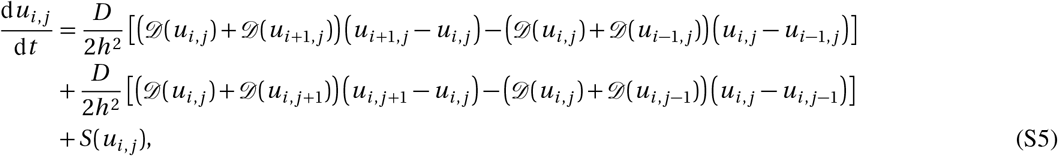

where we have approximated the internode diffusivity with an arithmetic average [72]. Equation S5 is valid at the central nodes, *i* = 1,2,…, *I* — 1 and *j* = 1,2,…, *J* — 1, and is modified on the boundary nodes since we implement symmetry (no-flux) boundary conditions along the boundaries where *i* = 1, *i = I*, *j* = 1 and *j = J* [72]. This system of (*I* + 1)^2^ coupled ordinary differential equations is then integrated through time using MATLABs ode45 routine [70].

## S4 Regression parameters in Figure 5

The parameters of the regression lines used to fit dimensionless briding time vs dimensionless initial pore size in the model in Figure 5 are listed in Table S4. The regression parameters of best fit *c* and *a* (slope and intercept, respectively) depend on the initial density *u*_0_ and threshold density *u_c_*. The last column lists the corresponding value of *Dλ* estimated using Eq. (7). For clarity, Figure 5 only shows results with *u_c_* = 0.2 and *u_c_* = 0.5.

**Table S4.** Regression parameters

| $u_0$ | $u_c$ | $c$ | $a$ | $D\lambda$ |
| --- | --- | --- | --- | --- |
| 0.2 | 0.2 | 0.69 | −0.15 | $184 \mu\text{m}^2/\text{day}^2$ |
| 0.5 | 0.2 | 0.66 | −0.79 | $167 \mu\text{m}^2/\text{day}^2$ |
| 0.8 | 0.2 | 0.63 | −1.02 | $153 \mu\text{m}^2/\text{day}^2$ |
| 0.2 | 0.5 | 0.67 | 0.26 | $176 \mu\text{m}^2/\text{day}^2$ |
| 0.5 | 0.5 | 0.66 | −0.59 | $172 \mu\text{m}^2/\text{day}^2$ |
| 0.8 | 0.5 | 0.64 | −0.92 | $164 \mu\text{m}^2/\text{day}^2$ |
| 0.2 | 0.8 | 0.63 | 1.18 | $154 \mu\text{m}^2/\text{day}^2$ |
| 0.5 | 0.8 | 0.66 | −0.03 | $173 \mu\text{m}^2/\text{day}^2$ |
| 0.8 | 0.8 | 0.68 | −0.68 | $183 \mu\text{m}^2/\text{day}^2$ |

## Footnotes

1 The average curvature is independent of any other pore shape characteristics. This is a consequence of the total curvature theorem which states that the total curvature of a closed curve in two-dimensional space is a topological invariant given by 2π times the turning number [28].

